# The P1 and P2 helices of the Guanidinium-II riboswitch interact in a ligand-dependent manner

**DOI:** 10.1101/2021.04.25.440196

**Authors:** Christin Fuks, Sebastian Falkner, Nadine Schwierz, Martin Hengesbach

## Abstract

Riboswitch RNAs regulate gene expression by conformational changes induced by environmental conditions and specific ligand binding. The guanidine-II riboswitch is proposed to bind the small molecule guanidinium and to subsequently form a kissing loop interaction between the P1 and P2 hairpins. While an interaction was shown for isolated hairpins in crystallization and EPR experiments, an intrastrand kissing loop formation has not been demonstrated. Here, we report the first evidence of this interaction *in cis* in a ligand and Mg^2+^ dependent manner. Using single-molecule FRET spectroscopy and detailed structural information from coarse-grained simulations, we observe and characterize three interconvertible states representing an open and kissing loop conformation as well as a novel Mg^2+^ dependent state for the guanidine-II riboswitch from *E. coli*. The results further substantiate the proposed switching mechanism and provide detailed insight into the regulation mechanism for the guanidine-II riboswitch class. Combining single molecule experiments and coarse-grained simulations therefore provides a promising perspective in resolving the conformational changes induced by environmental conditions and to yield molecular insights into RNA regulation.

## INTRODUCTION

Riboswitches are *cis*-regulatory elements that are located in the 5’ UTR of bacterial mRNA, affecting the expression of the downstream gene. They are generally comprised of an aptamer domain and an expression platform. The aptamer domain is responsible for specific ligand binding, whereas part of the expression platform can form distinct structure that trigger the genetic decision. By structurally coupling the aptamer domain with the expression platform, riboswitches are able to execute their regulatory function. There is a wide spectrum of metabolites that can be bound by the respective aptamer domain, ranging from ions (1,2), amino acids (3), nucleotides (4) and cofactors (5) to large biomolecules such as tRNAs (6). In general, two functional types of riboswitches can be discriminated: the transcriptional and translational riboswitches. In transcriptional riboswitches the expression platform contains a terminator sequence that halts transcription upon correct folding. In contrast, translational riboswitches contain an anti-Shine-Dalgarno (SD) sequence (7). Sequestering of the SD sequence prevents ribosome binding and thus translation initiation. In both cases ligand binding to the aptamer could either switch gene expression on or off, based on the type of riboswitch. This positive or negative feedback loop allows utilization of specific biosynthetic pathways (such as in the 2’dG riboswitch), or elimination of toxic substances, such as the fluoride riboswitch.

So far, four classes of riboswitches have been identified that bind the cationic molecule guanidinium (Gdm^+^): guanidine-I (8), -II (9), -III(2) and -IV (10). The corresponding genes are in most cases involved in Gdm^+^ detoxification, and code for proteins like guanidine-carboxylases or multidrug efflux pumps (e. g. SugE). The guanidine-II riboswitch is the smallest representative of guanidine riboswitches classes, and was named mini-*ykkC* prior to Gdm^+^ being identified as the ligand in 2017 (9). It consists of two GC-rich hairpins termed P1 and P2, both containing a conserved ACGR loop motif. It has been proposed that this class features a translational regulation mechanism since the two hairpins are connected with a linker containing a putative anti-SD sequence(11). So far, in-line probing experiments (9) and crystallization (12,11,13) analyses have suggested a kissing loop formation through canonical CG base pairs upon Gdm^+^ binding to the loop region (Figure 1A). This interaction supposedly sequesters the anti-SD sequence, exposing the SD sequence and thus facilitating translation of the downstream gene. Crystallization could however only show homo-dimerization of isolated hairpins. Recently, using EPR it was shown that RNAs comprising either hairpins P1 or P2 could form homo-as well as heterodimers (14).

**Figure 1:**
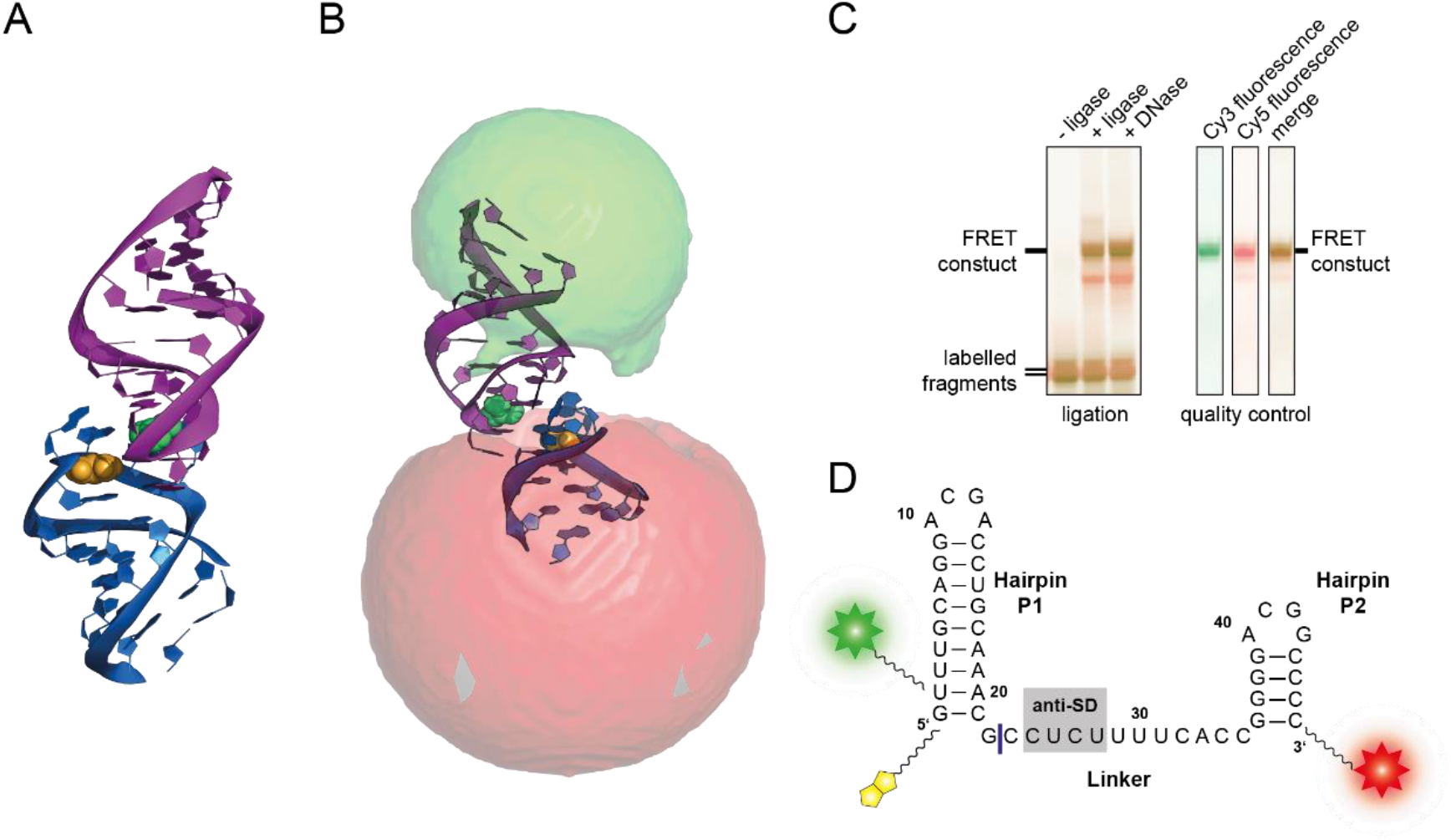
A) Crystal structure of *E. coli* SugE guanidine-II riboswitch P1 homodimer (PDB ID: 5NDI) (12) B) Accessible volume of the fluorophores calculated with FPS. One stem from the crystal structure was shortened according to the four base pairs in P2. C) Ligation reaction of the fluorophore labelled 23mer and 24mer fragments to yield the FRET construct (left) and PAGE analysis of the dual fluorescently labelled, purified construct (right). D) Sequence of the FRET construct used in this work. The biotin moiety is illustrated in yellow and the Cy3 and Cy5 attachments sites are shown as a green and red star, respectively. The ligation site between the 23mer and the 24mer is shown in blue.

Until now, there was however no direct evidence that an interaction in *cis* between the two hairpins can form in a functional RNA (14) and that this interaction is regulated by ligand binding. To fill this gap, we combined single-molecule FRET (smFRET) spectroscopy and coarse-grained simulations. The results from our complementary approach show that the riboswitch aptamer domain can adopt three different conformational states, including a ligand dependent state that involves intrastrand kissing loop formation.

## MATERIAL AND METHODS

### Construct design, dye attachment, and FRET positioning and screening software (FPS)

For this work the sequence of the *E. coli* SugE guanidine-II riboswitch aptamer was used (Figure 1D). The construct contained the P1 and P2 hairpin as well as the native linker connecting both hairpins. The 5’ C1 was exchanged with a G to stabilize the hairpin. Labelling sites were chosen at U3 via a C5 amino-allyl modification and the 3’ phosphate with a C6 amino-modifier. For immobilization, a biotin modifier was used at the 5’ end. The sequence was split into a 23mer and 24mer to allow separate labelling.

For FRET efficiency prediction, the FRET positioning and screening software (FPS) (15) was used. The crystal structure of the *E. coli* P1 hairpin homodimer (PDB: 5NDI (12)) was used as a template. One monomer was shortened to four basepairs in the stem to simulate the correct length of P2. For Cy3, dye radii of 6.8 Å, 3.0 Å, and 1.5 Å, respectively, were used. Cy5 was calculated with 11.0 Å, 3.0 Å, and 1.5 Å, respectively. The linkers were described with 22.3 Å and 4.5 Å for 5’ modification (C5 of amino-allyl uridine), and 27.1 Å and 4.5 Å for the 3’ modification (3’ oxygen). The Förster radius for the Cy3/Cy5 pair was set to 60 Å (16). The program was used to calculate the accessible volume clouds as depicted in Figure 1B, as well as the expected FRET efficiencies and average distances between the dyes.

### RNA synthesis: labelling

Modified RNAs for the FRET construct were purchased in two fragments (Dharmacon) (5’ fragment: Biotin-GU(5-NH2-U) UGC AGG ACG ACC UGC AAA CG, 3’ fragment: P-CCU CUU UUC ACC GGG GAC GGC CCC C6NH2). 30 nmol of each RNA were ethanol precipitated, and subsequently resuspended in 20 µL freshly prepared 0.1 M NaHCO_3_ (pH 8.0). Cy3 or Cy5 amine-reactive dyes (Amersham CyDye Mono-Reactive Dye Packs, GE Healthcare) were dissolved in 20 µL DMSO. Labelling was achieved by mixing the two solutions and incubation of the RNA with the respective dye for 90 min (3’ fragment) or 3 h (5’ fragment) at room temperature under light protection. RNA was precipitated and dissolved in 300 µL deprotection buffer (100 mM AcOH adjusted to pH 3.8 with TEMED) and incubated at 60 °C for 40 min (3’ fragment) or 2 h (5’ fragment). Deprotected RNA was precipitated, and non-biotinylated RNA was dissolved in 0.1 M TEAA (pH 7.0) and purified via reverse phase chromatography with an Äkta Basic system using a C8 column (Kromasil 100 C8 7µm 250×4.6mm). A gradient from 100% TEAA buffer to 50% MeCN was applied. Fractions with labelled RNA were collected and precipitated.

### RNA synthesis: ligation and purification

The FRET construct was synthesized through splinted ligation of the two fluorophore labelled fragments. All nucleic acid components had a final concentration of 10 µM each. RNA fragments were dissolved in water, heated to 95 °C for 2 min and placed on ice. The DNA splint (TGG GGC CGT CCC CGG TGA AAA GAG GCG TTT GCA GGT CGT CCT GCA AAC CTA TAG TGA GTC GTA TTA) and T4 ligase buffer (50 mM TRIS/Cl, 10 mM MgCl_2_, 1 mM ATP, 10 mM DTT, pH 7.5) were added and the mixture incubated at 85 °C for 3 min. After slowly cooling down T4 DNA ligase (final concentration of 40 U/mL, NEB) was added and the reaction was performed for 2 h at room temperature. 1 U Turbo DNase (Invitrogen) was added and incubated for 30 min at 37 °C, and subsequently extracted using phenol/ether extraction and precipitated. The FRET construct containing the desired sequence (Figure 1C) was separated via denaturing PAGE, eluted and ethanol precipitated.

### smFRET measurements

Microscope slides (quartz) and cover slips were cleaned with nitrogen plasma for 10 min. Channels were made by aligning parafilm stripes on the slide, covering it with the coverslip and heating everything to 80 °C for 30 s. Cooled down channels were filled with 1 mg/mL biotin labelled bovine serum albumin (BSA, Sigma-Aldrich) in T50 buffer (10 mM Tris/Cl, 50 mM NaCl, pH 8.0) and incubated for 2 min. Channels were then washed with 50 µL of T50 before incubation with 0.2 mg/mL streptavidin in T50 for 2 min. Channels were washed with 50 mM Tris/Cl (pH 7.4) with the respective Mg^2+^ and Gdm^+^ concentration. 100 pM RNA was folded in the respective buffer by incubation at 95 °C for 2 min and cooling on ice for 5 min, unless stated otherwise. RNA was flushed into the channel for immobilization. Prior to the measurement, the channel was rinsed with imaging buffer (sample conditions in Tris buffer, 10% (w/v) D-(+)-glucose, 80 µg/mL glucose oxidase, 20 µg/mL catalase, and Trolox (saturated)).

An objective-type spinning-spot total internal reflection microscopy setup with an EMCCD camera (iXon, Andor Technology) with 532 nm laser (green laser) and 633 nm laser (red laser) excitation with an integration time of 100 ms at 22 °C was used for smFRET measurements. For histograms 20 frames with green excitation were recorded. For kinetic data and verification of single step photobleaching movies of up to 7 min were recorded.

FRET efficiencies for histograms were calculated as the average over 2 s. FRET efficiencies were binned to a bin size of 0.05. The donor only peak was fitted with a Gaussian fit, and subtracted from the data. The remaining data was plotted into histograms. For these histograms, 3 states were fitted with a Gaussian fit using OriginPro 2018b (Northampton). The fractions of the individual states were calculated from the ratios of the area under the individual curves. For kinetic data, traces before photobleaching were manually selected for consistency, anticorrelated dye behaviour, and single-step photobleaching. The selected traces were stitched to a single trace of 50,000 datapoints, and Hidden Markov modelling software (HaMMy) (17) was applied using a 3 state model. Fitting the traces with a 5 state model did not result in additional, discernible FRET states (data not shown). Dwell times for each transition were fitted with an exponential decay and rate constants (k) were calculated using OriginPro 2018b.

The distribution of individual states of the dynamic molecules was calculated based on the rate constants k (Table 1 & 2) assuming the 3 state model shown in Figure 5C. For the equilibrium concentrations, the integrated phenomenological rate equations were used starting with equimolar concentrations (t=0) and iterating until an equilibrium was reached.

**Table 1:** Transition rates derived from dwell time fits at various Mg^2+^ concentrations. In cases where a reliable fit could not be obtained, i.e., due to an insufficient number of transitions (n), the rates were not determined (n.d.)

| Transition | 0 mM | 0.5 mM | 1 mM | 3 mM | 5 mM |
| --- | --- | --- | --- | --- | --- |
| $k_{U,K} [\text{s}^{-1}]$ | $5.6 \pm 0.14$<br>(n = 313) | $5.8 \pm 0.16$<br>(n = 626) | $3.2 \pm 0.04$<br>(n = 637) | $4.6 \pm 0.08$<br>(n = 541) | $4.0 \pm 0.07$<br>(n = 366) |
| $k_{K,U} [\text{s}^{-1}]$ | $1.1 \pm 0.03$<br>(n = 291) | $1.0 \pm 0.03$<br>(n = 575) | $1.6 \pm 0.03$<br>(n = 511) | $1.5 \pm 0.03$<br>(n = 439) | $2.2 \pm 0.11$<br>(n = 336) |
| $k_{U,M} [\text{s}^{-1}]$ | n.d.<br>(n = 21) | $0.8 \pm 0.04$<br>(n = 423) | $1.6 \pm 0.04$<br>(n = 339) | $2.0 \pm 0.07$<br>(n = 221) | $3.6 \pm 0.10$<br>(n = 110) |
| $k_{M,U} [\text{s}^{-1}]$ | n.d.<br>(n = 43) | $0.8 \pm 0.04$<br>(n = 476) | $0.6 \pm 0.03$<br>(n = 466) | $0.8 \pm 0.05$<br>(n = 324) | $0.8 \pm 0.10$<br>(n = 140) |
| $k_{K,M} [\text{s}^{-1}]$ | $0.7 \pm 0.05$<br>(n = 73) | $0.6 \pm 0.03$<br>(n = 168) | $0.9 \pm 0.05$<br>(n = 367) | $1.4 \pm 0.05$<br>(n = 315) | $2.2 \pm 0.07$<br>(n = 272) |
| $k_{M,K} [\text{s}^{-1}]$ | n.d.<br>(n = 51) | $1.2 \pm 0.03$<br>(n = 115) | $0.7 \pm 0.04$<br>(n = 240) | $0.4 \pm 0.03$<br>(n = 212) | $0.6 \pm 0.04$<br>(n = 243) |

**Figure 5.**
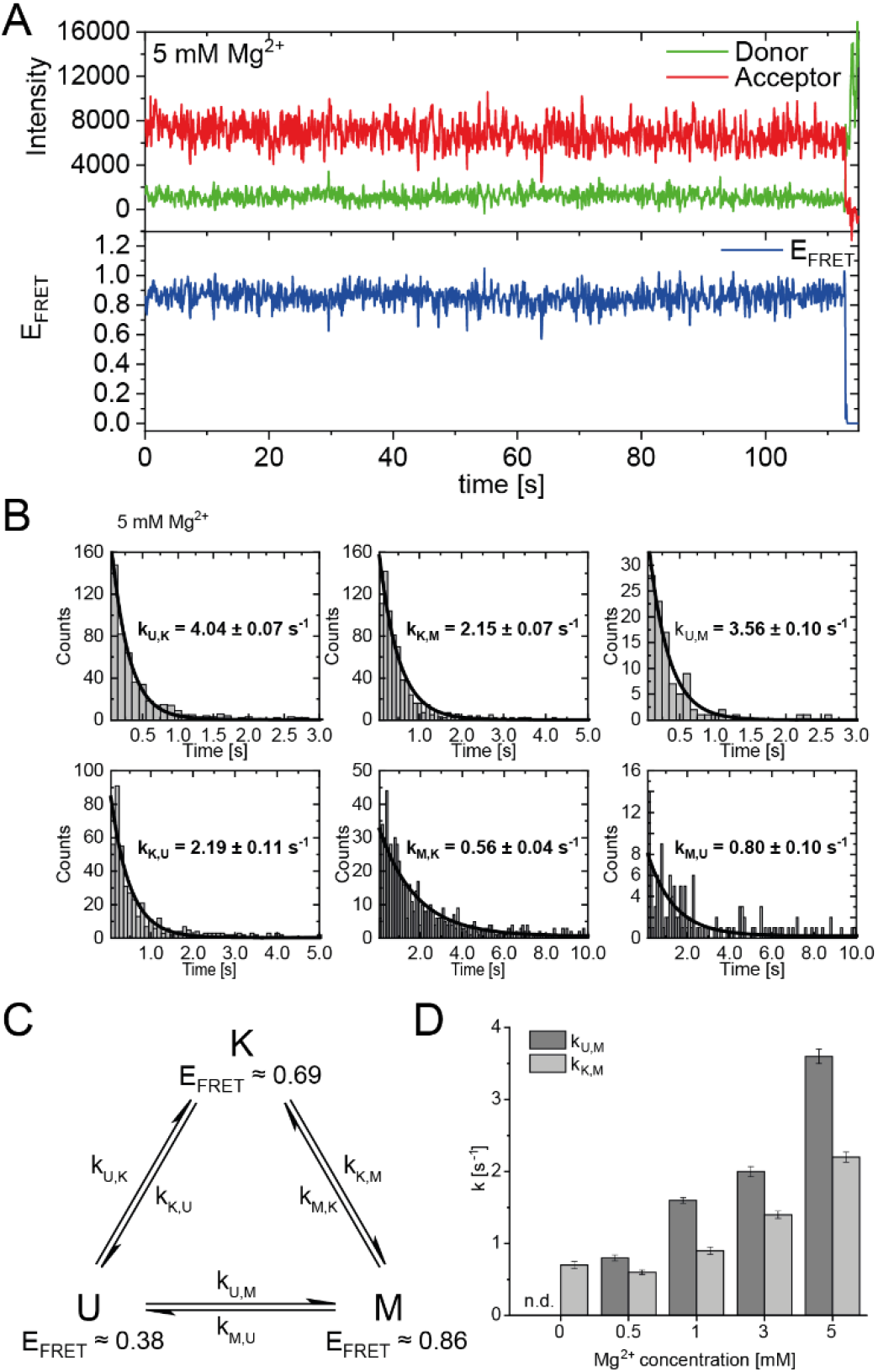
A) Exemplary FRET trace of a static molecule in the M-state. B) Exemplary dwell time plots used to determine transition rates. smFRET trajectories for each condition were fitted for 3 states with HaMMy and resulting dwell times were fitted with a monoexponantial decay. C) 3 state kinetic model D) Change of transition rates with increasing Mg^2+^ concentration of the transitions of the U to M (dark grey) and K to M (light grey).

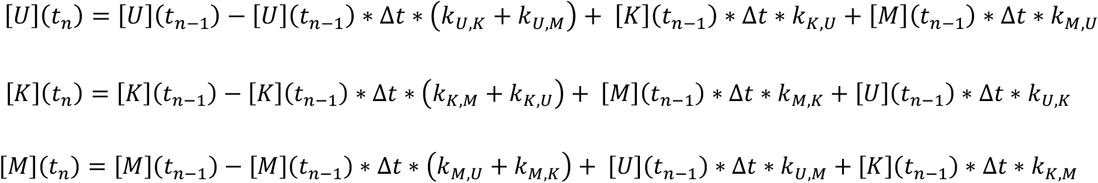

Here, [*i*](*t*_*n*_) are the concentrations of states i=U,K,M at time *t*_*n*_, k_i,j_ are the rate constants between states j and j and Δ*t* is the timestep used (0.1 s).

In the absence of Mg^2+^, the equilibrium concentrations were not calculated due to the missing rate constants k_U,M_, k_M,U_, and k_M,K_. For 1 mM Mg^2+^ and 30 mM Gdm^+^, transitions between the K- and M-state were considered to be 0.

### Coarse-grained simulations

The simulation construct was modeled based on a crystal structure of the *E. coli* guanidine-II riboswitch P1 stem-loop dimer (PDB: 5NDI) (12). One loop was shortened to the length of the wild type P2 stem-loop. Base pairs were mutated to match the FRET construct using Chimera (18). The linker between the P1 and P2 stem-loops was modeled using ModeRNA (19). The coarse-grained RNA simulations were performed using a three-interaction site model (TIS) developed by Thirumalai and coworkers (20–24). For our present study, TIS is particularly suited since it is computational efficient allowing us the investigation of folding/unfolding transitions while reproducing the folding thermodynamics with good accuracy (21). In the TIS model, the intramolecular attractive interactions are defined based on the residues that appear in the native structure. This description, inherent to all Gō-like models, ensures that the native structure is the minimum energy structure (25). Native hydrogen bonds and tertiary stacks in the simulation construct were defined based on the crystal structure of the P1 stem-loop dimer (12).

In order to capture folding intermediates that are not stabilized by native interactions, non-native secondary structure interactions were included via the base-stacking interactions of consecutive nucleotides and hydrogen-bond interactions between all nucleobases (22). A non-interacting adenine was added capping the structure at the 3’-end. The RNA was placed in a cubic box with an edge length of 70 nm. Simulations were run in the low friction regime for 1.5 × 10^9^ steps with a timestep of 0.05τ (τ = 50 fs) at 298.15 K. The cut-off for electrostatic interactions was set to 3 nm. Magnesium was included explicitly via the effective Magnesium-phosphate interaction potential derived from RISM theory (24). Simulations were performed with concentrations ranging from 0 to 10 mM. Monovalent ions were included implicitly at a concentration of 50 mM. To compare the simulations to experiments, the FRET efficiency was calculated. Structures from the simulations where backmapped to an atomistic representation. Accessible volumes of the FRET dyes and corresponding mean FRET efficiencies were estimated using *avtraj* (26). Dye parameters were matched to the previously stated FPS parameters. The resulting FRET efficiencies were subsequently binned using the same bin size (0.05) as in the experiments and fitted with the same routine described above. For comparison with experimental data the most probable FRET efficiency of each state is obtained from the maximum of the fitted Gaussian.

## RESULTS

### Construct design and synthesis

Several crystal structures have shown the formation of a kissing loop interaction involving canonical C-G base pairs between isolated hairpins (11–13), resulting in homodimeric structures. However, two problems can be envisioned when transferring these findings to functional constructs. First, the two stems in all natural riboswitches have different sequences. Secondly, the two hairpins are linked with a presumably regulatory anti-Shine-Dalgarno sequence. It is therefore unclear whether the interactions found in the crystal structures can actually be transferred to the natural riboswitch. Therefore, a linked construct comprised of the native hairpins is required to resolve whether the interaction between P1 and P2 helices can occur *in cis*.

Based on the available structural information from crystal structures (12) as well as in-line probing experiments (9), an RNA FRET construct was designed. For this, a 47mer RNA sequence of the guanidine-II riboswitch aptamer upstream of the *E. coli sugE* gene was selected (Figure 1C). Fluorophore attachment sites were placed in helical regions of both hairpins expecting them to assume a defined distance upon kissing loop formation, supposedly creating a signature FRET efficiency. A native U of the P1 stem was modified at position C5, and the second modification was introduced at the 3’ phosphate.

In order to identify the expected FRET efficiency of this construct in a potential kissing loop conformation (such as the one in the crystal structure), we modelled the available conformational states for the dyes with the FPS software package. This revealed a FRET efficiency of 0.72 (Figure 1B). Based on these results the RNA fragments were designed, and the construct successfully synthesized. We therefore divided the natural aptamer domain into two fragments corresponding to the 23mer P1 hairpin and the 24mer linker and P2 region (Figure 1C). The 5’ fragment had an additional conjugated biotin to allow immobilization required for smFRET measurements. Fragments were successfully labelled with Cy3 and Cy5 using NHS-chemistry, respectively. The final FRET construct was synthesized via subsequent splinted ligation of the dye-labelled fragments and denaturing PAGE purification of the double labelled RNA (Figure 1D).

### Initial smFRET characterization

Synthesis of this FRET construct now enables investigation whether P1 and P2 interact in *cis* using smFRET. Initially, we tested the FRET efficiencies and their response to selected Mg^2+^ ion and Gdm^+^ ligand conditions. In the absence of both ligand and Mg^2+^ the majority of molecules resided in a low FRET state (E_FRET_ ≈ 0.38) (Figure 2, top left). This low FRET state corresponds to a distance between the dyes that is larger than what would be expected for the conformation modelled from the crystal structure. As shown in the following, this state corresponds to an unfolded conformation, which we term the U-state.

**Figure 2:**
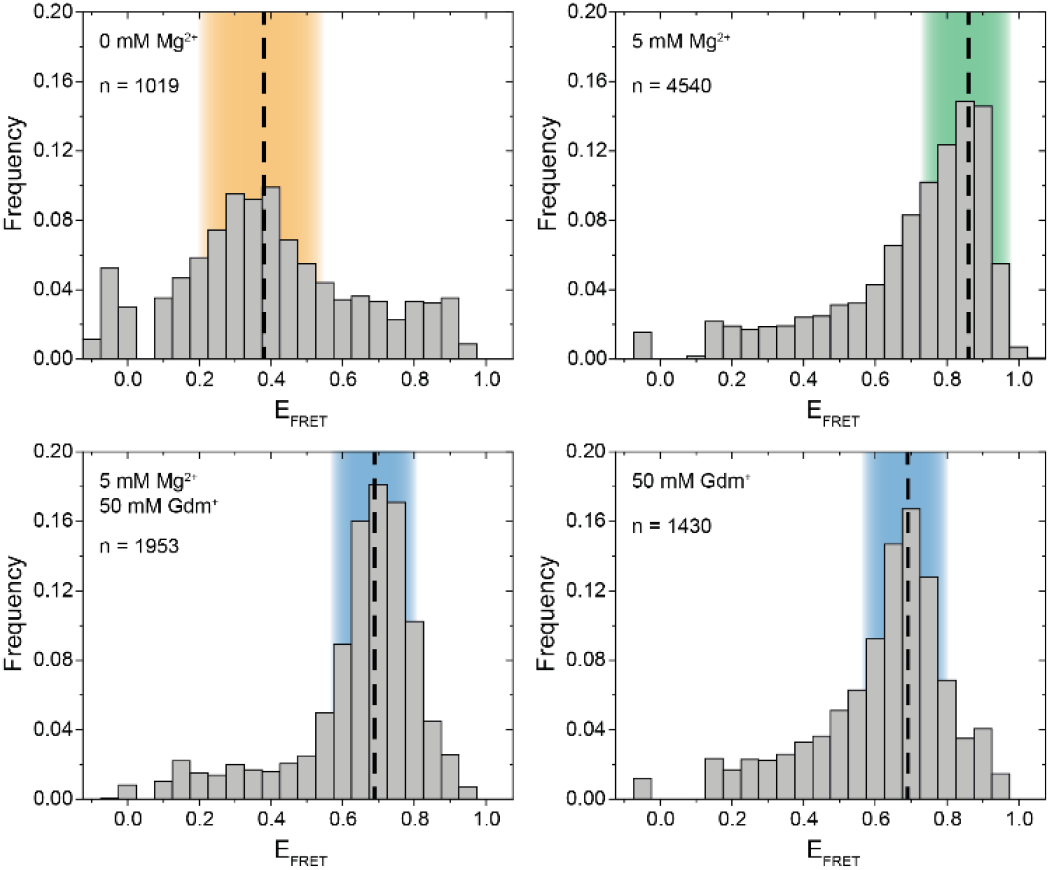
Initial smFRET experiments. The histograms show the populations at different FRET efficiencies in absence and presence of Mg^2+^ and Gdm^+^. The U-state is highlighted in orange, the K-state in blue and the M-state in green.

In presence of high Mg^2+^ concentrations, the riboswitch folds into a high FRET state (E_FRET_ ≈ 0.86) which we designate the M-state (Figure 2, top right).. The M-state has a higher FRET efficiency than expected from the crystal structure.

Upon addition of Gdm^+^ ligand, the equilibrium was shifted to an intermediate FRET efficiency of 0.69 (Figure 2, bottom). This FRET value is in excellent agreement with the FPS modelled distance in the crystal structure of single hairpins in a kissing loop interaction. Therefore, the intermediate FRET conformation is assigned as the K-state.

To summarize these initial experiments depending on the solution conditions, three distinct FRET states for the guanidine-II riboswitch were identified. For all of these states we verified that the data indeed originated from individual molecules by monitoring single step photobleaching in time resolved experiments. This strongly suggests that an RNA fold with fluorophore distances as expected for a kissing loop interaction between P1 and P2 is formed in *cis* within a full-length aptamer construct. Further, a previously uncharacterized M-conformation was identified.

For these experiments, the molecules were folded with the same conditions that were used for the subsequent smFRET measurements, which in principle also applies to data from in-line probing. The folding conditions for our experiments as well as those used for structural analysis of individual riboswitch hairpins however raise the question whether these states are in equilibrium or are conformationally trapped. Along the same line, for the function of some riboswitches it is crucial that the RNA does not only fold into different conformations, but that those are interconvertible in response to changing environmental conditions.

We therefore evaluated the reversibility of the RNA conformations by exchanging the buffer directly on the smFRET slide. FRET histograms in Figure S1 show that molecules on the same slide were able to shift from the U-state in absence of Mg^2+^ and Gdm^+^ to the M-state upon addition of Mg^2+^ ions. Upon further addition of Gdm^+^ the K-state was the predominant population, and subsequent removal of both Mg^2+^ and Gdm^+^ ions resulted in a quantitative return of the molecules to the low FRET U-state. This shows that the majority of molecules in our smFRET experiments retains responsiveness towards the buffer conditions and does not enter a conformationally trapped state.

### Coarse-grained simulations and smFRET analysis of Mg^2+^-dependent folding

We investigated the response of the riboswitch to changes in the Mg^2+^ concentration by performing Mg^2+^ titration experiments while monitoring the abundance of each of the three states. We found that high Mg^2+^ concentrations lead to stabilization of the M-state in smFRET experiments, while intermediate Mg^2+^ concentrations cause a shift from the U-to the K-state (Figure 3A & B, Figure S2A).

**Figure 3:**
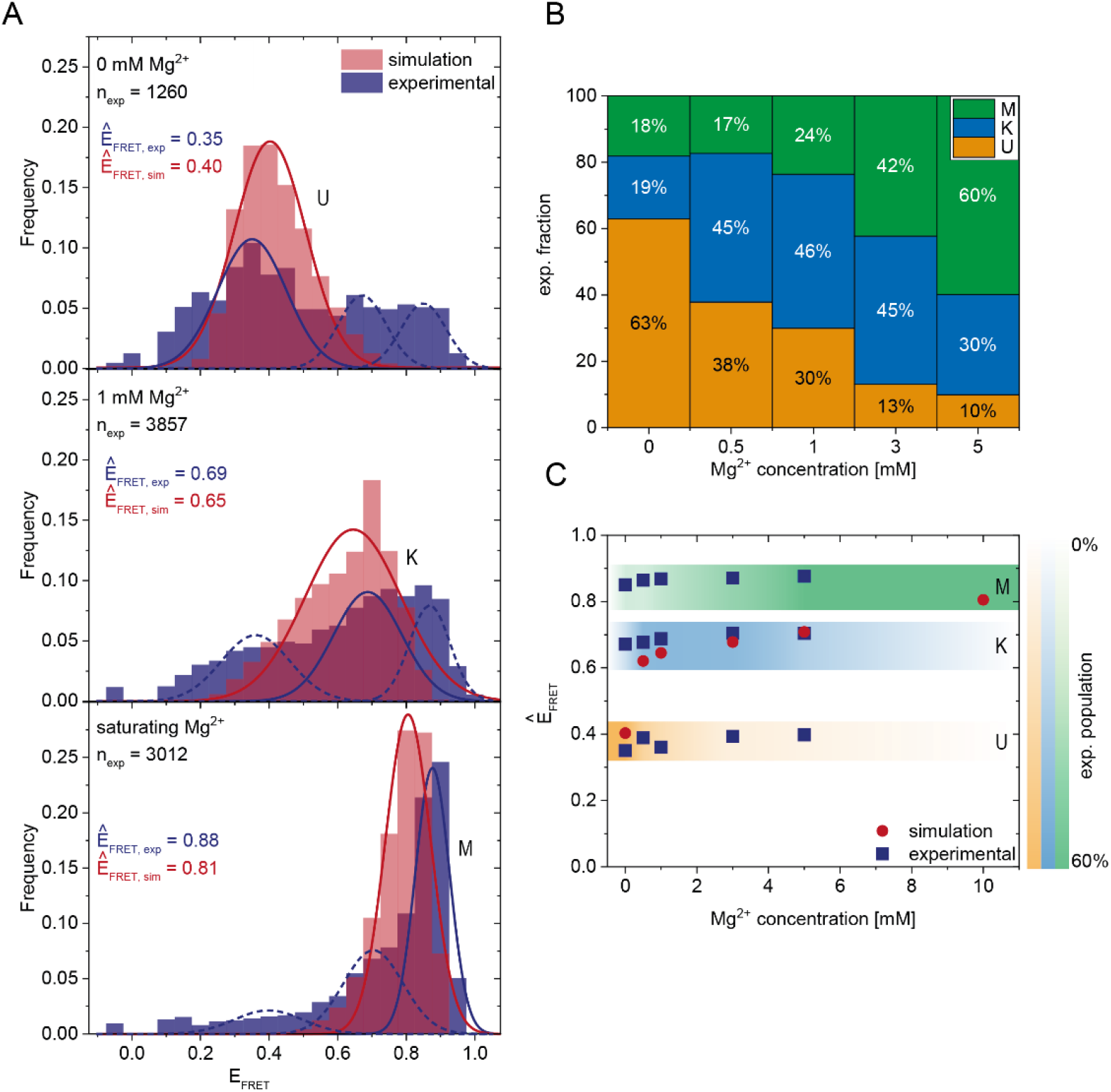
Comparison of experimental smFRET and simulated response of the riboswitch RNA to varying Mg^2+^ concentrations A) Histograms of the populations obtained in the smFRET experiments (blue) and in the coarse-grained simulations (pink) at different Mg^2+^ concentrations. The experiments were fitted with 3 Gaussians corresponding to the 3 states. The simulations were fitted with 1 Gaussian since at each concentration predominantly one state was populated while the probability of the other states was negligibly small. The maximum of the Gaussian fits Ê_FRET,exp_ and Ê_FRET,sim_ corresponds to the FRET efficiency of the most probable structure of each state and is shown as inset. B) Experimental fraction of riboswitches in the three states at various Mg^2+^ concentrations. The fractions were calculated from the area under the fits (shown in A). C) Comparison of most probable FRET efficiency Ê_FRET_ (corresponding to the maxima in the Gaussian fits shown in A) from experiments and simulations at various Mg^2+^ concentrations.

To further characterize these states, we performed coarse-grained simulations. In agreement with the experiments, the coarse-grained simulations suggest the existence of three different states dependent on the Mg^2+^ concentration. Representative structures of each state were selected from the simulations (Figure 4). At low Mg^2+^ concentrations, the linker in the obtained structures is extended. As a result, the P1 and P2 stem-loops are pointing in different directions, and the tetraloop sequences are not in spatial proximity. At intermediate Mg^2+^ concentrations, we find mainly structures with a native-like kissing loop orientation of the P1 and P2 stem-loops. At 10 mM Mg^2+^, more compact structures with a different secondary structure are most abundant. While the P1 hairpin remains folded, the P2 stem unfolds and forms new basepairs with the linker containing the putative anti-SD sequence (Figure 4).

**Figure 4:**
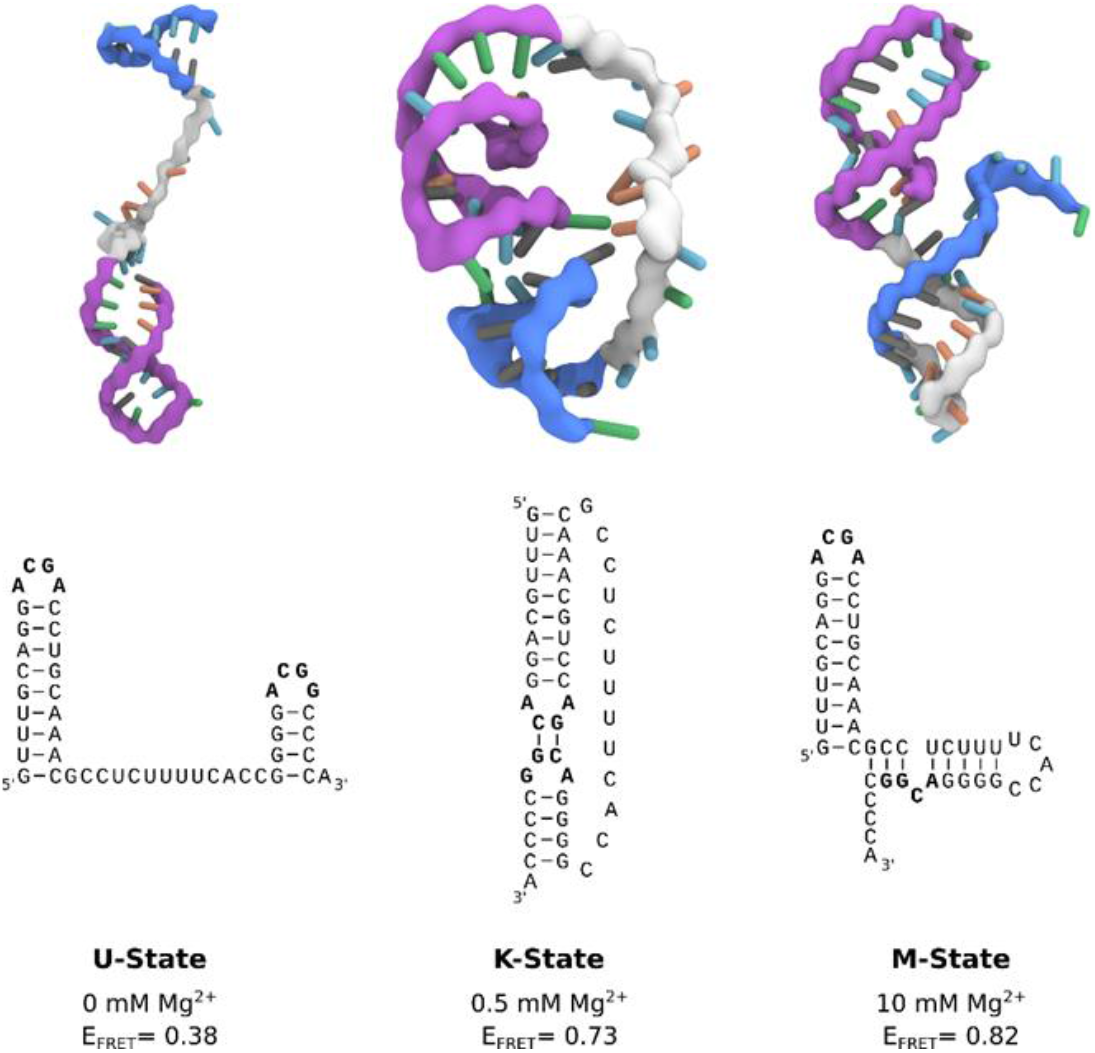
Representative structures of the three states from coarse-grained simulations at different Mg^2+^ concentrations. The P1 and P2 stem-loops are coloured in purple and blue respectively, the linker is coloured in grey. The corresponding secondary structure is shown below each three-dimensional structure. The calculated FRET efficiency (E_FRET_) of each structure averaged over different dye orientations is given.

To assess whether the conformations derived from the simulations and the FRET experiments are representatives of the same state, we calculated the theoretical FRET efficiencies and their distributions from the simulations. In the coarse-grained simulations, the native K-state is designed to be the minimum energy structure by including the native contacts of the kissing-loop interaction. At the same time, the coarse-grained model includes non-native interactions and therefore allows us to capture the U-state at low Mg^2+^ concentrations and the M-state at high concentrations. Due to the Gō-like nature of the model, the abundance of the conformations in each state does not correspond to an equilibrium distribution. In addition, we find predominantly one conformational state and interconversion between the states is rare. Therefore, the magnitude of the FRET frequency from simulations and experiments varies as expected (Figure 3A).

Still, the FRET efficiencies of the most probable structure in each state from simulations and experiments (Ê_FRET_) can be compared directly. Figure 3A, C show that Ê_FRET_ from experiments and simulations is in very good agreement over a range of Mg^2+^ concentrations. This in turn suggests that the structures derived from the coarse-grained simulations and the ones observed in the FRET experiments at different concentrations are in fact identical.

These results show that combining coarse-grained simulations and smFRET experiments provides detailed, complementary insights into the response of the riboswitch RNA to different Mg^2+^ concentrations. They also reveal that a coaxial orientation of the hairpins necessary for the kissing loop conformation is accessible in the absence of the ligand. High Mg^2+^ concentrations lead to the folding of the riboswitch into an alternative M-conformation. Here, the coarse-grained simulations allow us to resolve the alternative base pairing pattern and the three-dimensional structure.

### Analysis of dynamics: Mg^2+^ titration

After identification and structural description of the three states U, K and M, and showing that the states are interconvertible, we characterized the transitions between these states further. To this end, we performed time resolved smFRET measurements and observed the behavior of the molecules over several minutes at different Mg^2+^ concentrations in the absence of Gdm^+^. While some of the traces remained in one FRET state throughout the measurement (example in Figure 5A), the majority of molecules showed multiple transitions, in which each of the states was accessible directly from every other state. We isolated FRET data prior to photobleaching for each molecule and stitched these traces together to a single FRET trace up to 50,000 datapoints (Figure S3). From these stitched traces, we performed hidden Markov Modelling using the HaMMy software (17) with a three state model (Figure 5C). Since the number of transitions for each molecule was significantly higher than the number of stitched molecules, we were able to extract kinetic information (Table *1*) about the fast-transitioning molecules (in other cases we note n.d.). For this we plotted the dwell times and fitted a mono-exponential function to derive rate constants (Figure 5B).

We found that the transition from the U-state to the K-state was fast, with a rate constant of approximately k_U,K_ = 4.6 s^−1^ (average over all Mg^2+^ concentrations). We note that this transition is a reorientation of the two hairpins P1 and P2 rather than opening of any helix, as suggested by the structures from the coarse-grained simulations. For opening the kissing loop, we found rates of k_K,U_ between 1.0 and 2.2 s^−1^. For the formation of the M-conformation a likely Mg^2+^-dependent behaviour from both the U- and the K-state was observed, in which rates increased slightly with Mg^2+^ concentration (Figure 5D & Table *1*). This increase ranged from 0.8 to 3.6 s^−1^ for K_U,M_ and from 0.6 to 2.2s^−1^ for k_K,M_. Refolding into both the U- and K-state starting from the M-state occurred slower in comparison, with no obvious dependence on Mg^2+^ concentrations for K and U (around k_M,U_ = 0.8 s^−1^ and k_M,K_ = 0.7 s^−1^).

### Effect of Gdm^+^ ligand on structural dynamics

After observing the K-state to a certain amount (46% at 1 mM Mg^2+^) in the absence of the ligand and a stabilization of this conformation (to 66%) at high Gdm^+^ concentrations (Figure 2), we characterized the ability of P1 and P2 to form a kissing loop orientation in a ligand dependent manner. We chose the intermediate (and likely physiological) Mg^2+^ concentration of 1 mM to perform ligand titrations. Increasing Gdm^+^ concentrations in the sub-millimolar range did not result in significant shifts between the populations, with 30 to 41% of the molecules adopting the K state. Addition of 30 mM Gdm^+^ however shifted the populations into the K state (66%) on the expense of both U and M states, confirming a distinct ligand dependence of this conformation (Figure 6A&B & Figure S2C). The high concentrations of Gdm^+^ used in this experiment could however also have an unspecific chaotropic effect. We therefore repeated this measurement in the presence of 30 mM urea instead of Gdm^+^ to further assess the specificity of ligand binding. In contrast to Gdm^+^, high concentrations of urea did not shift the equilibrium towards the K-state, but rather resulted in a destabilization of the RNA as evidenced by an increased and likely broadened U-state population (35% of molecules) (Figure 6C).

**Figure 6:**
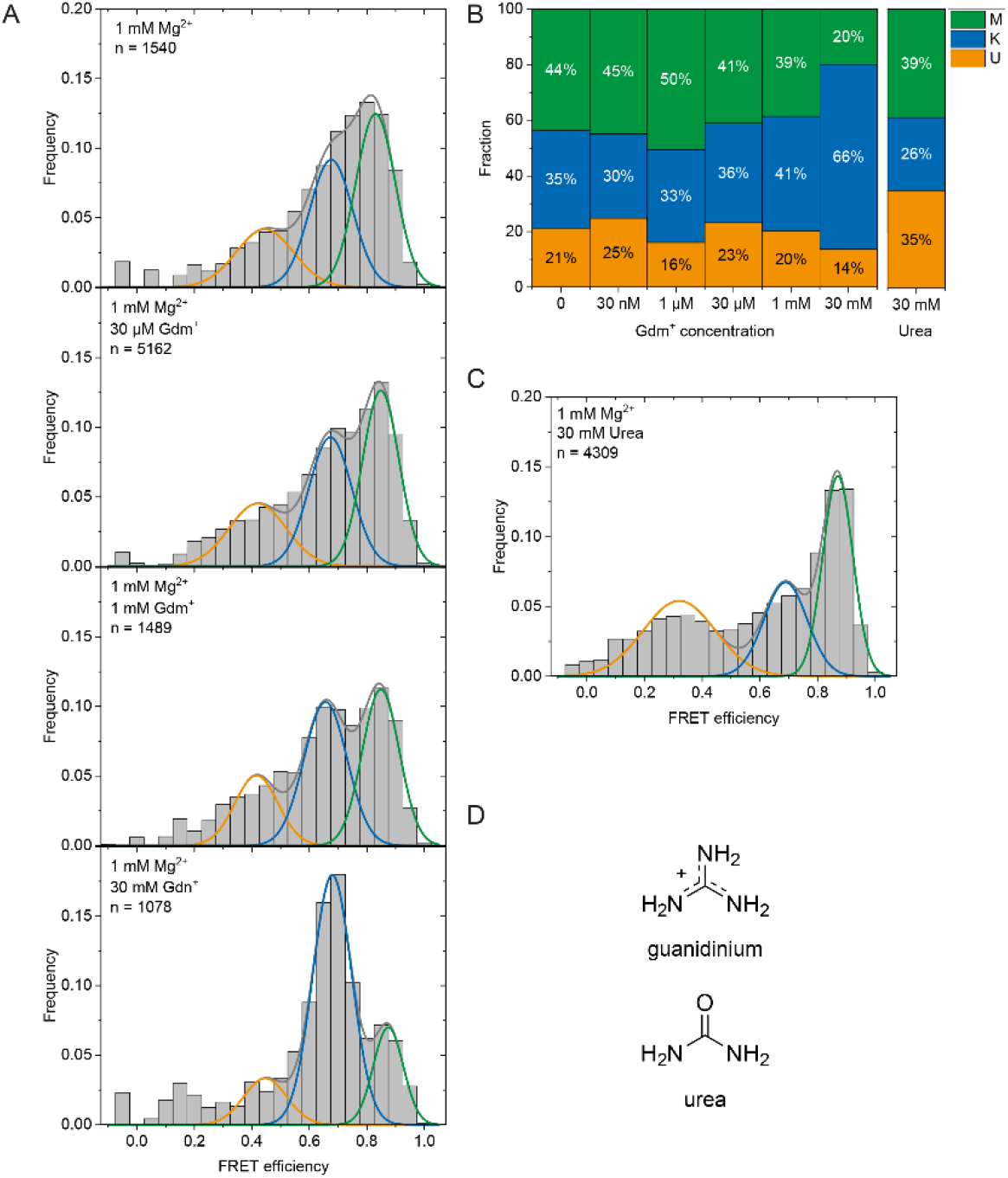
A) Histogram analysis of the Gdm^+^ titration at constant 1 mM Mg^2+^. The data was fitted with 3 Gaussian fits showing the change between the K-state (blue) and the M-state (green) with increasing Gdm^+^ concentration. B) Histogram analysis at 1 mM Mg^2+^ with 30 mM urea. C) Fractions of molecules adopting the individual states based on the area under the fitted curves. D) Structures of Gdm^+^ and urea.

As for the Mg^2+^ dependence we also calculated the rate constants for all observable state transitions in our Gdm^+^ titration (Figure 7B & Table 2) from the dwell times derived from stitched traces (Figure S4). Here, transitions between the U- and K-states are comparable with the data obtained from Mg^2+^ titration experiments, i.e. with k_U,K_ = 3.9 s^−1^ and k_K,U_ = 1.3 s^−1^, respectively. In general, all other observed transition rates were also comparable with the data of the Mg^2+^ titration in absence of ligand. Within error, we were not able to identify a faithful ligand dependence of any of the fast transitions, with the exception of k_U,K_, which upon increase of the ligand concentration rose from ≤3.7 s^−1^ up to 1 mM Gdm^+^ to 6.4 s^−1^ at 30 mM Gdm^+^.

**Table 2:** Transition rates derived from dwell time fits at various Gdm^+^ concentrations at 1 mM Mg^2+^. In cases where a reliable fit could not be obtained, i.e. due to a low number of transitions (n), the rates were not determined (n.d.)

| Transition | 0 | 30 nM | 1 μM | 30 μM | 1 mM | 30 mM |
| --- | --- | --- | --- | --- | --- | --- |
| $k_{U,K} [\text{s}^{-1}]$ | $3.2 \pm 0.04$<br>(n = 737) | $3.5 \pm 0.04$<br>(n = 730) | $2.9 \pm 0.04$<br>(n = 929) | $3.7 \pm 0.07$<br>(n = 934) | $3.7 \pm 0.04$<br>(n = 682) | $6.4 \pm 0.09$<br>(n = 933) |
| $k_{K,U} [\text{s}^{-1}]$ | $1.6 \pm 0.03$<br>(n = 729) | $1.0 \pm 0.05$<br>(n = 696) | $0.8 \pm 0.05$<br>(n = 903) | $1.9 \pm 0.03$<br>(n = 846) | $1.2 \pm 0.06$<br>(n = 632) | $1.4 \pm 0.03$<br>(n = 930) |
| $k_{U,M} [\text{s}^{-1}]$ | $1.6 \pm 0.04$<br>(n = 79) | $2.4 \pm 0.11$<br>(n = 95) | $1.5 \pm 0.05$<br>(n = 917) | $1.4 \pm 0.05$<br>(n = 215) | $3.3 \pm 0.09$<br>(n = 107) | $5.0 \pm 0.19$<br>(n = 49) |
| $k_{M,U} [\text{s}^{-1}]$ | $0.6 \pm 0.03$<br>(n = 86) | $0.3 \pm 0.04$<br>(n = 129) | $0.3 \pm 0.03$<br>(n = 223) | $0.4 \pm 0.03$<br>(n = 303) | $0.4 \pm 0.04$<br>(n = 157) | $0.6 \pm 0.11$<br>(n = 51) |
| $k_{K,M} [\text{s}^{-1}]$ | $0.9 \pm 0.05$<br>(n = 71) | $0.8 \pm 0.05$<br>(n = 143) | $0.7 \pm 0.06$<br>(n = 162) | $1.0 \pm 0.07$<br>(n = 428) | $1.5 \pm 0.08$<br>(n = 402) | n.d.<br>(n = 26) |
| $k_{M,K} [\text{s}^{-1}]$ | $0.7 \pm 0.04$<br>(n = 64) | $0.2 \pm 0.03$<br>(n = 109) | $0.2 \pm 0.02$<br>(n = 135) | $0.3 \pm 0.02$<br>(n = 340) | $0.4 \pm 0.02$<br>(n = 353) | n.d.<br>(n = 23) |

**Figure 7:**
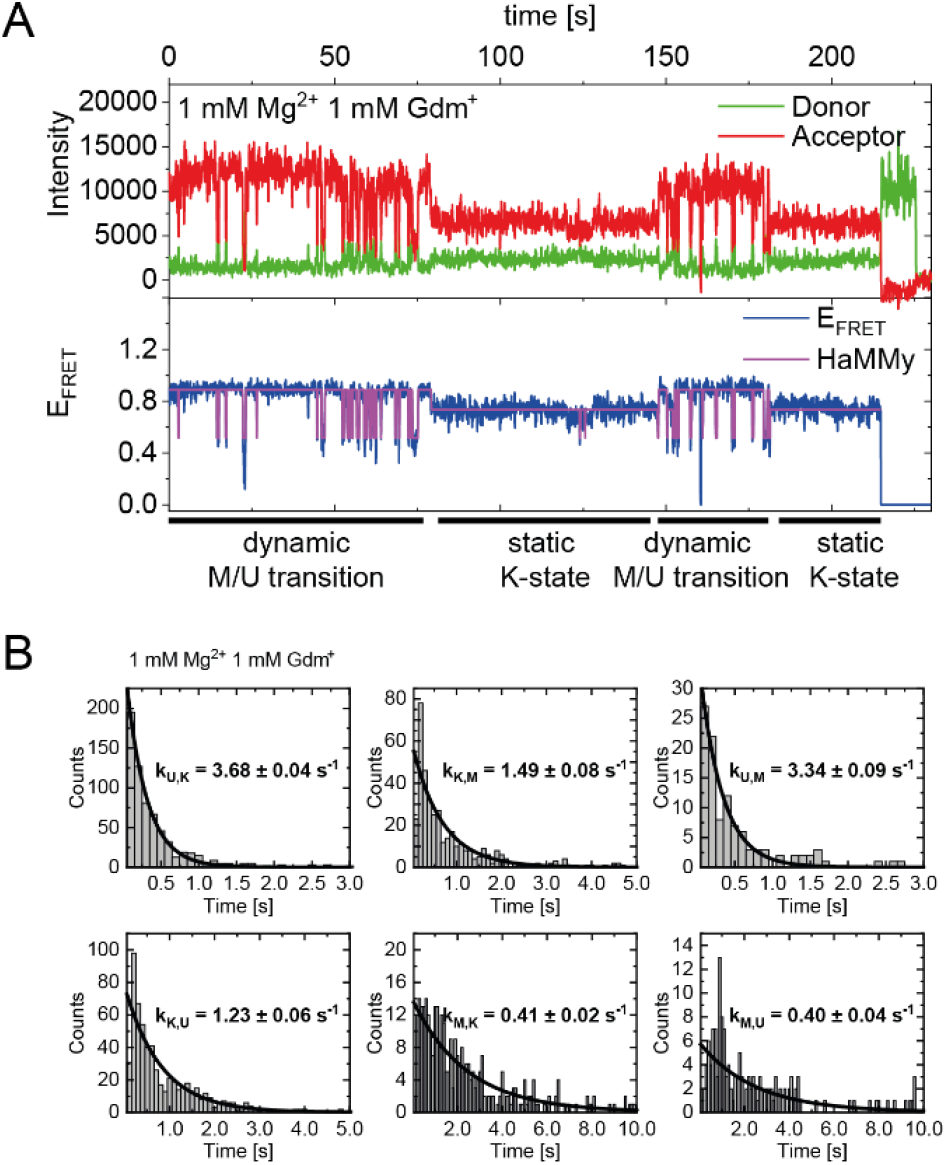
A) Exemplary FRET trace of a dynamic molecule switching between a dynamic M-state and a static K-state. B) Exemplary dwell time plots of 1 mM Mg^2+^ and ligand each, used for transition rate calculations.

As can be seen in Figure 7A, we do however observe some molecules that transiently adopt the K-state for an extended period of time (> ∼20sec). In addition to that, all of the data sets contained a limited number of molecules that adopted a particular state (U, K, or M) and showed no transitions between states for the duration of the observation (termed “static molecules”). Moreover, due to our limited observation time (minute timescale), the number of the transitions may not be faithfully represented in our steady state analysis shown in Figure 6. We therefore calculated the population distributions using the phenomenological equations described in the Materials and Methods section (Figure S5).When comparing the transition rates of the dynamic molecules (Tables 1 and 2) with the populations obtained from all molecules (i.e. Figure 1, Figure 6), we find that these are not always in agreement.

## DISCUSSION

Using retrosynthetic splitting, we successfully designed and synthesized a smFRET construct for analysis of individual guanidine-II riboswitch RNA aptamer domain molecules. This RNA showed a distinct and reversible response to binding of both Mg^2+^ and Gdm^+^ ions and folding into three discernible states (**U**nfolded, **K**issing loop orientation, and **M**g^2+^ dependent). Binding of the ligand Gdm^+^ resulted in a FRET efficiency that closely matched the values derived from FPS-based modelling of the crystal structure. Since our methodological approach can robustly identify signals from individual molecules, this is a very strong indication that the interaction between P1 and P2 helices can indeed occur *in cis*, resulting in a kissing loop structure.

The experimental results are supported further by coarse-grained simulations in the absence of Gdm^+^ ligand. From these simulations, we can identify and structurally characterize three distinct Mg^2+^-dependent states. At low Mg^2+^ concentrations, an unfolded structure with folded P1 and P2 hairpins is observed. This conformation is shifted to a kissing loop structure at intermediate concentrations, emphasizing the requirement for Mg^2+^ ions for correct folding of the RNA. Further increasing the Mg^2+^ concentration leads to a conformation that exhibits a distinctly different secondary structure and base pairing pattern for the shorter P2 hairpin. In summary, the results suggest that the RNA requires a certain window of Mg^2+^ concentrations that facilitate folding into the kissing loop structure. When comparing theoretical FRET values derived from simulations and experiments, we find very good agreement of the individual FRET efficiency values and remarkably consistent results of the response to varying Mg^2+^ concentrations. Interestingly, the structure resolved by the simulations at high Mg^2+^ concentrations is fully consistent with the measured FRET values even in absence of any other, orthogonal *a priori* structural knowledge. Furthermore, the structure is in agreement with other data from in-line probing experiments (9), which show a rather low cleavage intensity of the P2 loop even in absence of ligand, but at high (20 mM) Mg^2+^ concentrations. This may be a feature specific to the sequence of this riboswitch in *E. coli* but emphasizes the importance of a comparison of data points obtained at different Mg^2+^ concentrations. Since the state occurs at elevated Mg^2+^ concentrations and to a smaller extent also at near-physiological Mg^2+^ concentrations, an unambiguous judgment on a possible regulatory relevance of this state is not possible. As can be seen in Figure 3, the anti-SD-sequence would be sequestered in this conformation, since it is interacting with a part of the sequence that is otherwise involved in formation of the P2 hairpin.

In our smFRET analysis, we find that refolding from the U into the M state apparently increases with Mg^2+^ concentration over the physiologically relevant range. For ligand binding, an increase of folding from the U into the K state requires very high (30 mM) concentration of ligand. While the K_d_ for this RNA is generally high (300 µM, (9)), this points to an even higher concentration that is required for full stability of the kissing loop interaction. We would like to note that with our experimental smFRET methodology we cannot provide a detailed structural characterization of the kissing loop interaction. Nevertheless, the longevity of some molecules in the K-state suggests a certain degree of stabilization rather than a statistical orientation of the hairpins. Since we also observe the K state in absence of the native ligand Gdm^+^, we cannot definitively identify whether Gdm^+^ ligand binding induces the kissing loop interaction (“induced fit”), or whether binding of the ligand merely stabilizes molecules that have already adopted a kissing loop orientation (“conformational selection”).

In our smFRET analysis, we expectedly find molecules that show multiple transitions in each time trace, but also molecules that occupy the same state for the duration of the experiment (dynamic versus static molecules, respectively). At the same time, the distribution of the entirety of molecules (as shown in the histograms in Figures 2, 5, and 7) shows somewhat different distributions between the U, K, and M state than what can be derived from the rates between those states (Figure S5). This can be rationalized by the presence of such static molecules, since these would contribute to the histograms, but not to the analysis of transition rates.

With regard to the Mg^2+^ titration, a consistent picture emerges from our combined approach of smFRET experiments and coarse-grained simulations. While smFRET experiments are routinely employed to characterize the Mg^2+^-dependent folding of RNAs (27–31) and in particular riboswitches (32–35), the combined approach used here allows us to provide a more comprehensive view. The quantitative agreement between the maximum efficiencies of each state from simulations and experiments highlights the compatibility of both approaches despite possible limitations in the time scale accessible to the simulations. In particular, the simulations complement the experiments by providing molecular insight into the base pairing pattern and the three-dimensional structures at different Mg^2+^ concentrations.

The comparison of simulations and smFRET data at different Mg^2+^ concentrations and the presence of the M state also show that a near-physiological range of Mg^2+^ ion concentrations is required for the capacity for efficient folding into the suggested functional K state. The K state in turn is significantly stabilized by ligand binding, confirming the functional relevance of our analysis. This also for the **first time** experimentally directly demonstrates that the functionally relevant interaction between the two hairpins P1 and P2 occurs *in cis* in a ligand-dependent manner.

## Conclusion

In summary, smFRET analysis and simulations of the full-length guanidine-II riboswitch aptamer domain from *E*.*coli* shows that the RNA can adopt three distinct states which are responsive to ligand as well as to Mg^2+^ concentration. In close agreement between experiments and coarse-grained simulations, we find that three interconvertible states: An unfolded state, a novel Mg^2+^-dependent state, and the presumably functional kissing loop interaction state.

Our results show that combining coarse-grained simulations and single-molecule FRET experiments provides complementary and detailed insights into the conformational changes induced by environmental conditions.

In light of the proposed translation regulation properties of this riboswitch, our findings are in excellent agreement with existing data from other methodological approaches (9,11,12,14). Our results further provide the first direct evidence of the ligand-dependent kissing loop orientation in *cis* for the guanidine-II riboswitch.

## SUPPLEMENTARY DATA

Supplementary Data are enclosed as 1 PDF file.

## ACKNOWLEDGEMENT

The authors would like to thank Prof. Harald Schwalbe for critical discussion and constant support, Prof. Josef Wachtveitl, Dr. Anna Wacker and Tatjana Schamber for critical discussions, and Prof. Mike Heilemann for access to instrumentation. N.S. and S.F. thank Dave Thirumalai, Naoto Hori and Hung T. Nguyen for fruitful discussions and support with TIS. LOEWE CSC and GOETHE HLR are acknowledged for supercomputing access.

