## Supplementary information for "The P1 and P2 helices of the Guanidinium-II riboswitch interact in a ligand-dependent manner"

### Supplementary Figure 1:

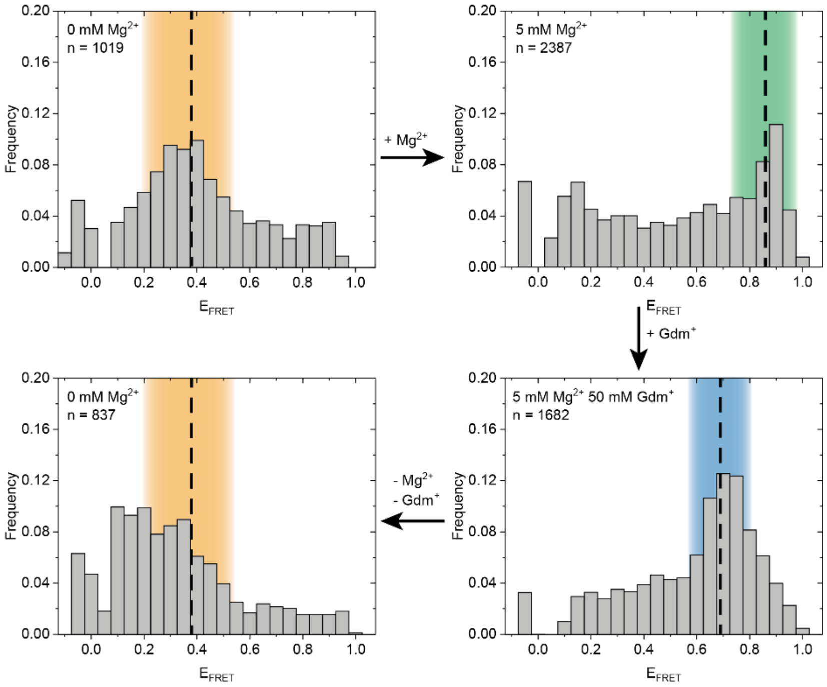


Figure S1: Buffer exchange smFRET experiments. The molecules were folded in absence of $Mg^{2+}$ and $Gdm^{+}$ ions and measured (upper left). Subsequently the buffer was exchanged to $Mg^{2+}$ containing buffer without heating steps and measured again (upper right). The buffer was exchanged again first to high concentrations of $Mg^{2+}$ and $Gdm^{+}$ (lower right) before returning to the starting buffer without $Mg^{2+}$ and $Gdm^{+}$ (lower left).

### Supplementary Figure 2:

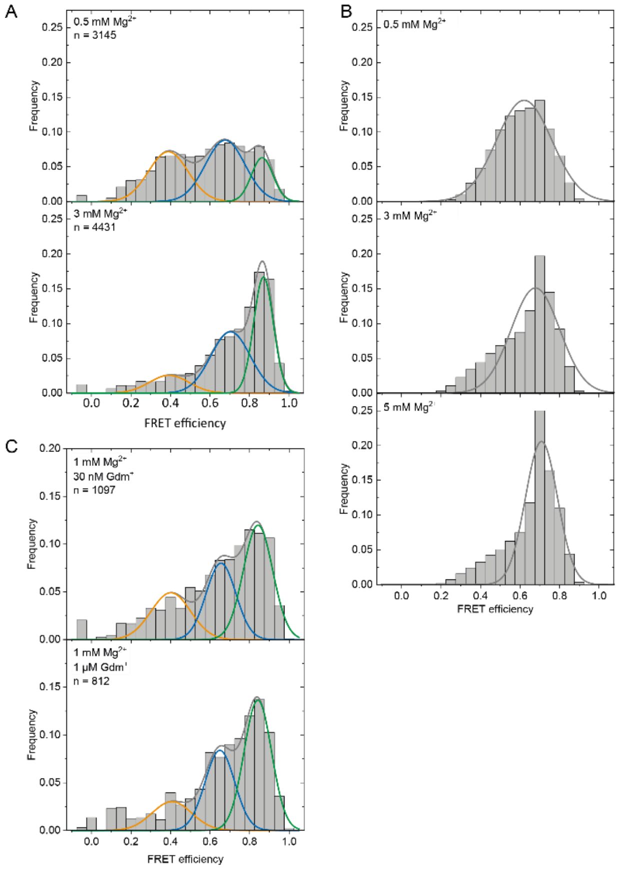


Figure S2: A) Additional experimental smFRET data from the $Mg^{2+}$ titration in absence of ligand fitted with 3 Gaussian fits. B) Additional histograms derived from MD simulation data from the $Mg^{2+}$ titration fitted with 1 Gaussian fit. C) Additional experimental smFRET data from the $Gdm^{+}$ titration at constant $Mg^{2+}$ concentration of 1 mM

### Supplementary Figure 3:

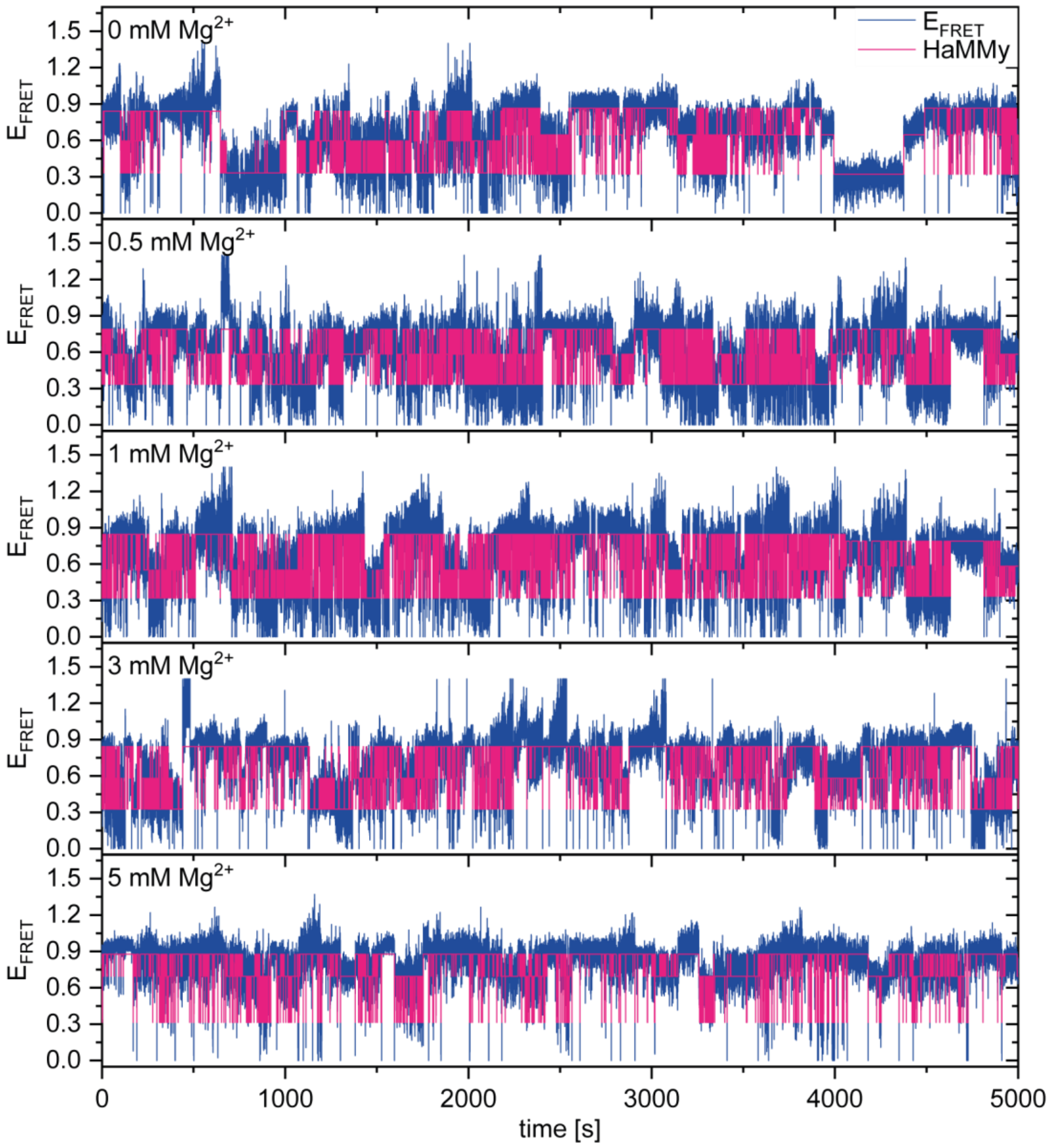


Figure S3: Stitched and HaMMy fitted smFRET traces from the $Mg^{2+}$ titration used for kinetic analysis. FRET trajectories were selected manually and stitched to a single trace with 50000 datapoints (blue). Each trace was subjected to a 3 state Hidden Markov modelling using HaMMy (pink).

**Supplementary Figure 4:**

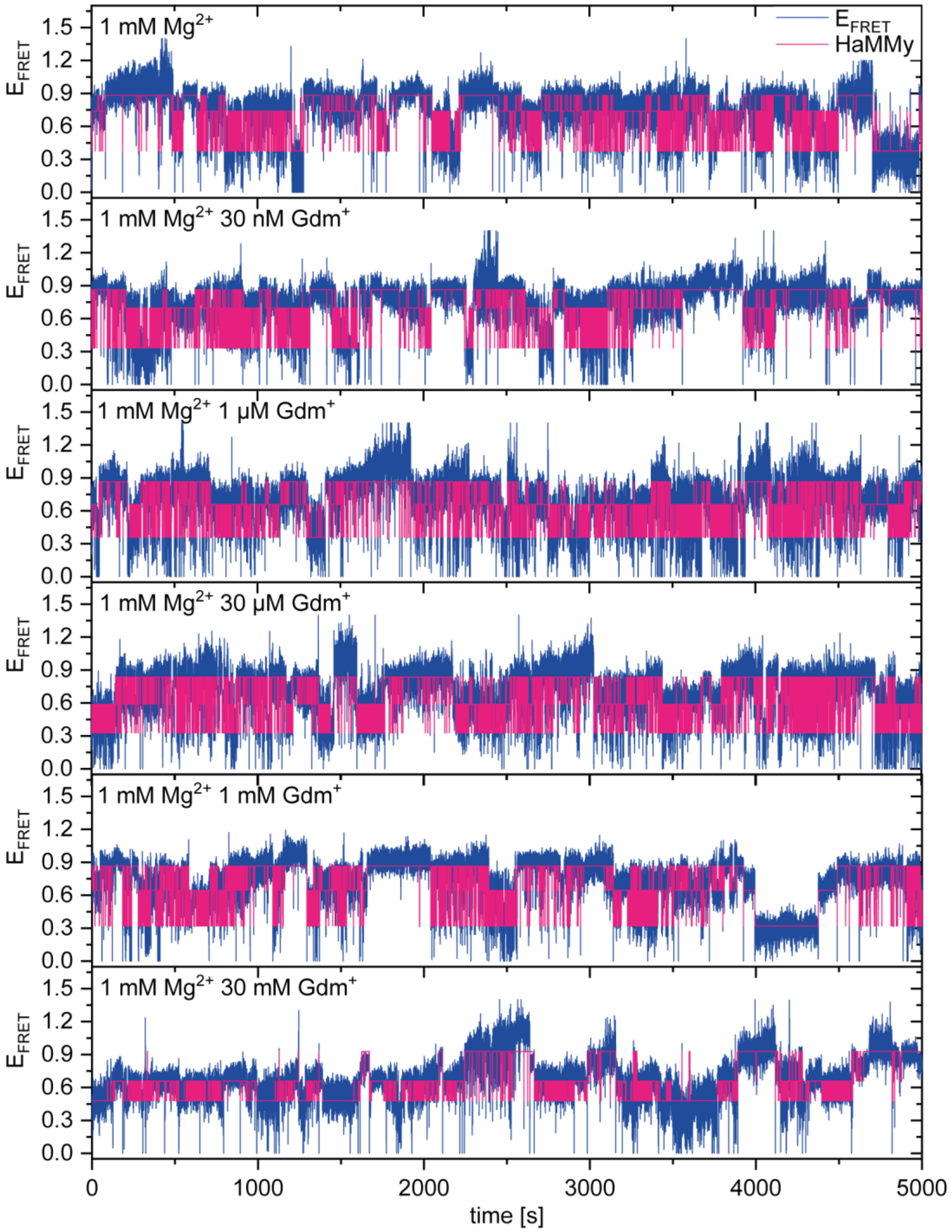


Figure S4: Stitched and HaMMy fitted smFRET traces from the $Gdm^{+}$ titration at constant 1 mM $Mg^{2+}$ used for kinetic analysis. FRET trajectories were selected manually and stitched to a single trace with 50000 datapoints (blue). Each trace was subjected to a 3 state Hidden Markov modelling using HaMMy (pink).

**Supplementary Figure 5:**

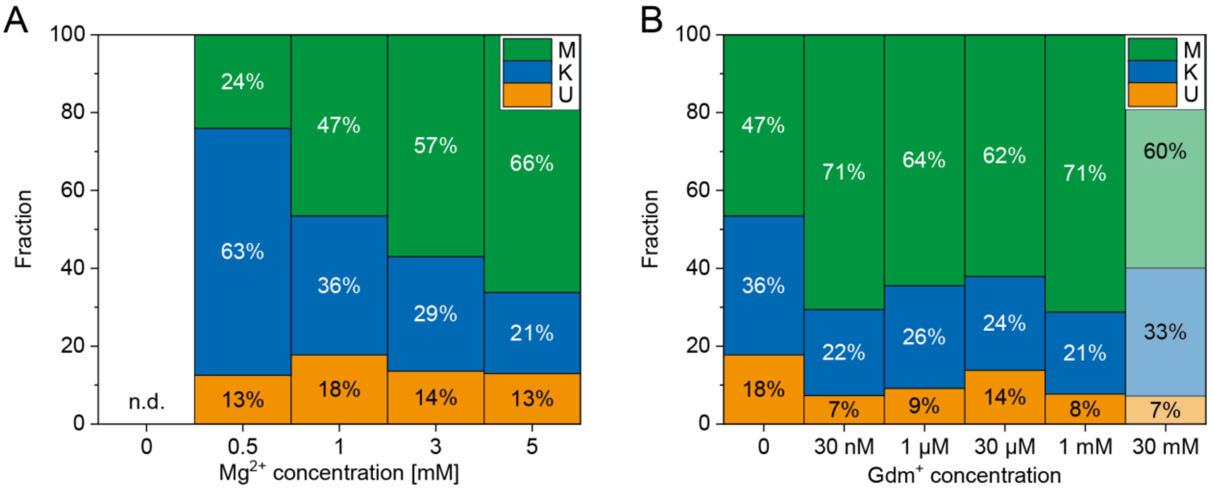


Figure S5: Distribution of individual states calculated from the rate constants of A) the $Mg^{2+}$ titration in the absence of $Gdm^{+}$ (Table 1) and B) the $Gdm^{+}$ titration at 1mM $Mg^{2+}$ (Table 2). Ratios for 30 mM $Gdm^{+}$ were calculated without transitions between K and M-state.
